# Antiviral CD19^+^CD27^+^ Memory B Cells Are Associated with Protection from Recurrent Asymptomatic Ocular Herpes Infection

**DOI:** 10.1101/2021.10.05.463293

**Authors:** Nisha R. Dhanushkodi, Swayam Prakash, Ruchi Srivastava, Pierre-Gregoire A. Coulon, Danielle Arellano, Rayomand V. Kapadia, Raian Fahim, Berfin Suzer, Leila Jamal, Lbachir BenMohamed

## Abstract

Reactivation of herpes simplex virus 1 (HSV-1) from latently infected neurons of the trigeminal ganglia (TG) leads to blinding recurrent herpetic disease in symptomatic (SYMP) individuals. Although the role of T cells in herpes immunity seen in asymptomatic (ASYMP) individuals is heavily explored, the role of B cells is less investigated. In the present study, we evaluated whether B cells are associated with protective immunity against recurrent ocular herpes. The frequencies of circulating HSV-specific memory B cells and of memory follicular helper T cells (CD4^+^ T_fh_ cells), that help B cells produce antibodies, were compared between HSV-1 infected SYMP and ASYMP individuals. The levels of IgG/IgA and neutralizing antibodies were compared in SYMP and ASYMP individuals. We found that: (*i*) the ASYMP individuals had increased frequencies of HSV-specific CD19^+^CD27^+^ memory B cells; and (*ii*) high frequencies of HSV-specific switched IgG^+^CD19^+^CD27^+^ memory B cells detected in ASYMP individuals were directly proportional to high frequencies of CD45R0^+^CXCR5^+^CD4^+^ memory T_fh_ cells. However, no differences were detected in the level of HSV-specific IgG/IgA antibodies in SYMP and ASYMP individuals. Using the UV-B-induced HSV-1 reactivation mouse model, we found increased frequencies of HSV-specific antibody-secreting plasma HSV-1 gD^+^CD138^+^ B cells within the TG and circulation of ASYMP mice compared to SYMP mice. In contrast, no significant differences in the frequencies of B cells were found in the cornea, spleen, and bone-marrow. Our findings suggest that circulating antibody-producing HSV-specific memory B cells recruited locally to the TG may contribute to protection from symptomatic recurrent ocular herpes.

**IMPORTANCE:** Reactivation of herpes simplex virus 1 (HSV-1) from latently infected neurons of the trigeminal ganglia (TG) leads to blinding recurrent herpetic disease in symptomatic (SYMP) individuals. Although the role of T cells in herpes immunity against blinding recurrent herpetic disease is heavily explored, the role of B cells is less investigated. In the present study, we found that in both asymptomatic (ASYMP) individuals and ASYMP mice there was increased frequencies of HSV-specific memory B cells that were directly proportional to high frequencies of memory T_fh_ cells. Moreover, following UV-B induce reactivation, we found increased frequencies of HSV-specific antibody-secreting plasma B cells within the TG and circulation of ASYMP mice, compared to SYMP mice. Our findings suggest that circulating antibody-producing HSV-specific memory B cells recruited locally to the TG may contribute to protection from recurrent ocular herpes.

## INTRODUCTION

With over a billion individuals worldwide currently infected with herpes simplex virus type 1 (HSV-1), herpes remains one of the most prevalent viral eye infections (1-3). Ocular herpes is mainly caused by HSV-1, which infects the cornea and then establishes latency in sensory neurons of the trigeminal ganglia (TG). Sporadic spontaneous reactivation of HSV-1 from latently infected neurons leads to viral shedding in saliva and tears which can ultimately cause symptomatic recurrent Herpes Stromal Keratitis (HSK), a blinding corneal disease. Despite the availability of many intervention strategies, the global picture for ocular herpes continues to deteriorate (4). Current anti-viral drug therapies (e.g., Acyclovir and derivatives) do not eliminate the virus and reduce recurrent herpetic disease by only ∼45% (5, 6). The challenge in developing an effective herpes treatment or vaccine is to determine the immune correlates of protection (7-13). Profiling humoral immunity in asymptomatic (ASYMP) and symptomatic (SYMP) HSV-1 infected individuals will further help in the 1) understanding if SYMP individuals have dampened humoral immune response in natural infection, 2) knowing immune correlates of protection that will help to understand vaccine efficacy, and 3) standardizing methods to explore vaccine efficacy.

Although the role of CD4^+^ and CD8^+^ T cells against HSV-1 reactivation is heavily explored, the role of B cells is less investigated. Unlike T cell memory which can be tissue-resident, recent studies suggest a migratory role for memory B cells (14, 15). Memory B cells (MBCs) are potential antibody secreting immune cells that differentiate following exposure to the virus. Following differentiation, MBCs remain in the peripheral circulation and secrete antibodies when they are re-exposed to their cognate antigen (16). MBCs in HSV-1 infected humans is not feasible to explore due to the low abundance of MBCs in peripheral blood. In fact, the low abundance of MBCs for any specific virus makes it challenging to study frequency, specificity, and breadth for the virus of interest. MBCs can be identified by their B-cell receptor (BCR), a membrane bound immunoglobulin (Ig) identical to the antibody they secrete upon activation (17, 18). In this study, we explored the frequency of MBCs found in the circulation of asymptomatic and symptomatic HSV-1 infected individuals by two different approaches –antigen-specific flow cytometry and ELISPOT.

To understand the mechanisms that cause a differential humoral response between SYMP and ASYMP herpes individuals, we studied the levels of several B cell associated ligands and cytokines like APRIL, BAFF, IL-21, IL-10, IL-17 and TNF-β in serum. APRIL (a proliferation inducing ligand) and BAFF (B cell activating factor) are ligands that can be released by proteolytic cleavage to form active, soluble homotrimers (19-21). They are expressed by immune cells other than B cells and their receptors BCMA (B cell maturation antigen) and TACI (TNFR homolog transmembrane activator and Ca^2+^ modulator and CAML interactor), are expressed exclusively by B cell lineage (22-24). IL-21, apart from its regulatory function plays a role in maintenance of antigen-specific memory B cells (25) and IL-10 is known to be involved in germinal center B cell responses (26).

Using the UV-B-induced HSV-1 reactivation mouse model, the frequencies of HSV-1 specific memory B and plasma cells were compared between SYMP and ASYMP mice by flow cytometry and ELISpot within cornea, TG, spleen and bone marrow and circulation. We found a significant increase in the frequencies of HSV-specific antibody-secreting plasma B cells within the TG of ASYMP mice compared to SYMP mice. In contrast, no significant differences were found in the cornea, spleen and bone-marrow. Our findings suggest that circulating HSV-specific antibody-producing memory B cells recruited locally at the TG site could contribute to protection from symptomatic recurrent ocular herpes.

## MATERIALS AND METHODS

### Human study population and PBMC isolation

Twenty-two volunteers (10 ASYMP, 10 SYMP and 2 HSV-1& HSV-2 negative) were enrolled in this study (**Table 1**). Peripheral blood was collected and PBMCs (peripheral blood mononuclear cells) were isolated by Ficol-Paque density gradient. The cells were then washed in PBS and re-suspended in complete culture medium consisting of RPMI-1640 medium containing 10% FBS (Bio-Products, Woodland, CA) supplemented with 1x penicillin/L-glutamine/streptomycin, 1x sodium pyruvate, 1x non-essential amino acids. The blood plasma was stored in -20°C for evaluation of herpes-specific antibodies later. HSV-1 infection was confirmed using a commercially available kit that detects anti-gG antigen of HSV-1 (HerpeSelect1 IgG ELISA, Focus Diagnostics, Cypress, CA).

### Flow cytometry

PBMCs of HSV-1 infected individuals were stained *ex-vivo* with the following flow antibodies listed below. Additionally, HSV-1 gD antigen (Virusys Corporation, Taneytown, MD) was coupled using FITC or AP lightning kit (Novus Biologicals, Littleton, CO) to detect HSV-1 specific B memory cells *ex-vivo* (27). ASYMP and SYMP mice were euthanized and immune cells from peripheral blood, spleen, bone marrow, TG and cornea were collected for flow cytometry staining for memory B cells and T follicular helper (Tfh) cells. Harvested TG were digested with collagenase III (5mg/ml) in RPMI 1640 containing 10% fetal bovine serum (FBS), 1% antibiotic/antimycotic, and gentamicin at 37° C. TG and cornea were dissociated with a 3-mL syringe-plunger head in the presence of media. Cell suspensions were passed through a 40-micron filter before staining. Single cell suspensions were labeled with the following fluorochrome-conjugated monoclonal antibodies: anti-mouse B220, CD73, PD-L2, CD80, CD45(A20), CD3(145-2C11), CD4(GK1.5), CXCR5, CD44, CD62L, ICOS, PD-1 (BD Biosciences, San Jose, CA). For surface staining, mAbs were added against various cell markers to a total of 1 ×10^6^ cells in phosphate-buffered saline (PBS) containing 1% FBS and 0.1% sodium azide (fluorescence-activated cell sorter [FACS] buffer) and left for 45 minutes at 4°C. For intracellular/intranuclear staining, cells were first treated with cytofix/cytoperm (BD Biosciences) for 30 minutes. Upon washing with Perm/Wash buffer, mAbs were added to the cells and incubated for 45 minutes on ice in the dark, washed with Perm/TFFACS buffer and fixed in PBS containing 2% paraformaldehyde. Labeled cells were suspended in 1% BSA in PBS and analyzed using the BD Fortessa flow cytometer. Intracellular staining was performed to detect HSV-1 specific plasma cells.

### HSV-1 gD -specific ASC ELISPOT assay

PBMCs from human or immune cells were stimulated (2-4 million cells/ ml) in B-cell media containing Human or mouse polyclonal B cell activator (Immunospot) for 5 days. Both CTL Human B-Poly-S and CTL Mouse B-Poly-S are stock solutions containing Resiquimod and either recombinant Human IL-2 or recombinant Mouse IL-2 respectively, used for the polyclonal expansion of memory B cells. This will activate the memory B cells to ASC. Cells were then washed in RPMI medium and plated in specified cell numbers in ELISPOT membrane plates coated with either HSV-1 gD antigen (Virusys) (1ng/well) or IgG/ IgA capture antibody (Immunospot Basic ELIPOT kits). The ASC secreting cells were detected after 48 hours of addition of cells to ELISPOT plates.

### HSV-1-specific IgA/IgG ELISA assay

Sera or plasma was isolated from blood by centrifugation for 10 min at 800g. Heat-inactivated HSV-1 McKrae strain was used for coating ELISA plates (Nunc Immunosorbent). The affinity of binding of antigen-specific antibodies to the HSV-1 McKrae strain was measured by ELISA plates coated overnight at 4°C with (10^4^ pfu/ well) heat-inactivated HSV-1. Heat-inactivated human or mouse serum (56°C for 1 hour) was then incubated at a specified dilution overnight at 4°C. HSV-1 specific IgG / IGA were then detected using human or mouse IgG/ IgA detection antibodies conjugated to HRP. Subsequently, TMB substrate was added to stop the reaction before performing the reading at 450 nm in the ELISA plate reader (iMark Microplate Reader, Bio-Rad).

### Viral plaque neutralization assay

Neutralizing antibody titers were determined by incubating 100 PFU of HSV-1 strain McKrae with serial dilutions of serum starting at 1:40 for 1 hour at 37°C. The endpoint neutralization titer was determined by the plaque assay on RS cells and was calculated as the serum dilution that reduced the number of plaques by 50% compared with PBS controls.

### Virus propagation and titration

For virus propagation, rabbit skin (RS) cells (ATCC, Manassas, VA) were grown in Minimum Essential Medium Eagle with Earl’s salts and L-Glutamine (Corning, Manassas, VA) supplemented with 10% fetal bovine serum and 1% penicillin-streptomycin. The HSV-1 laboratory strain McKrae was propagated in RS cells and purified by ultracentrifugation in sucrose gradient and titrated by the plaque assay.

### Mice and infection

All animals were handled with care according to the guidelines of American Association for Laboratory Animal Science (AALAS). For primary herpes infection, six-to eight-week-old male and female B6 mice were purchased from the Jackson Laboratory. The mice were anaesthetized with xylazine (6.6mg/kg) and ketamine (100mg/kg) prior to infection. Both corneas in each mouse was briefly scarified with a 25-gauge needle, tear film blotted, and 1×10^6^ pfu/eye of HSV-1 (strain McKrae) in 2 μL of sterile PBS were inoculated into the cornea. Corneal infection in all the infected mice was confirmed by viral plaque assay in tear swabs. Virus shedding in tear swabs was collected at day 2, 7 and 7 post-infections (p.i.). At day 35 p.i., eyes were reactivated by exposure to UV-B radiation for one minute and at day 6 post-reactivation, mice were categorized into ASYMP or SYMP depending on disease occurrence. ASYMP and SYMP mice were euthanized and immune cells from peripheral blood, spleen, bone marrow and TG were collected for flow cytometry staining for memory B cells and Tfh cells.

### Quantification of infectious virus

Tears were collected from both eyes using a Dacron swab (type 1; Spectrum Laboratories, Los Angeles, CA) on days 3, 5 and 7 p.i. Individual swabs were transferred to a 2mL sterile cryogenic vial containing 1ml culture medium and stored at -80°C until further use. The HSV-1 titers in tear samples were determined by standard plaque assays on RS cells as previously described (28). Eye swabs (tears) were analyzed for viral titers using the plaque assay. RS cells were grown to 70% confluency in 24-well plates. Infected monolayers were incubated at 37°C for 1 hour and rocked every 15 minutes for viral adsorption and then overlaid with medium containing carboxymethyl cellulose. After 48 hours of incubation at 37°C, cells were fixed and stained with crystal violet, and viral plaques and counted under a light microscope. Positive control assays used previously titrated laboratory stocks of McKrae.

### B cell development by Luminex

Asymptomatic and symptomatic patients’ serum (heat-inactivated at 56°C for 30 minutes) were assayed for cytokines involved in B cell development namely APRIL, BAFF, IL-10, IL-21, IL-7 and TNF-β using the Luminex kit according to the manufacturer’s instructions (R & D systems). Samples were assayed using the Luminex assay system (Magpix).

### Statistical analyses

Data for each assay were compared by analysis of variance (ANOVA) and Student’s *t* test using GraphPad Prism version 5 (La Jolla, CA). Differences between the groups were identified by ANOVA and multiple comparison procedures, as we previously described (29). Data are expressed as the mean ± SD. Results were considered statistically significant at *p* < 0.05.

## RESULTS

### 1. Increased frequency of HSV-1 specific circulating memory B cell in ASYMP HSV-1 infected individuals

Asymptomatic and symptomatic HSV-1 infected individuals were recruited to the study to understand the role of B cells in herpes. We collected their peripheral blood samples. PBMCs from ASYMP and SYMP individuals were stained for total B cells (CD19^+^), memory B cells (CD19^+^CD27^+^), IgG memory B cells (IgG^+^CD19^+^CD27^+^), IgA memory B cells (IgA^+^CD19^+^CD27^+^), IgM memory B cells (IgM^+^CD19^+^CD27^+^). ASYMP and SYMP individuals showed no differences in these general B cell profiles (**Fig. S1**). We stained the ASYMP and SYMP individuals for HSV-1 gD antigen specific memory B cells. HSV-1 specific memory B cells were studied by both antigen specific flow cytometry and ELISA. PBMC *ex vivo* was stained for CD19^+^ CD27^+^ B cells and analyzed for HSV-1 gD antigen binding cells. As expected, the percentage of herpes specific memory B cells were very low in the peripheral blood. Memory B cells express the antibody on their surface. HSV-1 gD antigen was conjugated to Alexa-fluor 488 and A647 fluorophores. Cells binding to both the antigen bound fluorophore were gated as HSV-1 specific cells. To confirm the gate, we performed florescence minus one (FMO) (**Fig. 1A**). We found an increased percentage of HSV-1 gD binding memory B cells in circulation of ASYMP individuals compared to SYMP herpes positive subjects [0.639 ± 0.102 % Vs 0.358± 0.070 %] (**Fig. 1B**). HSV-1 and HSV-2 negative individuals showed no detectable memory B cells compared to ASYMP individuals and SYMP herpes positive subjects (data not shown).

**Figure 1:**
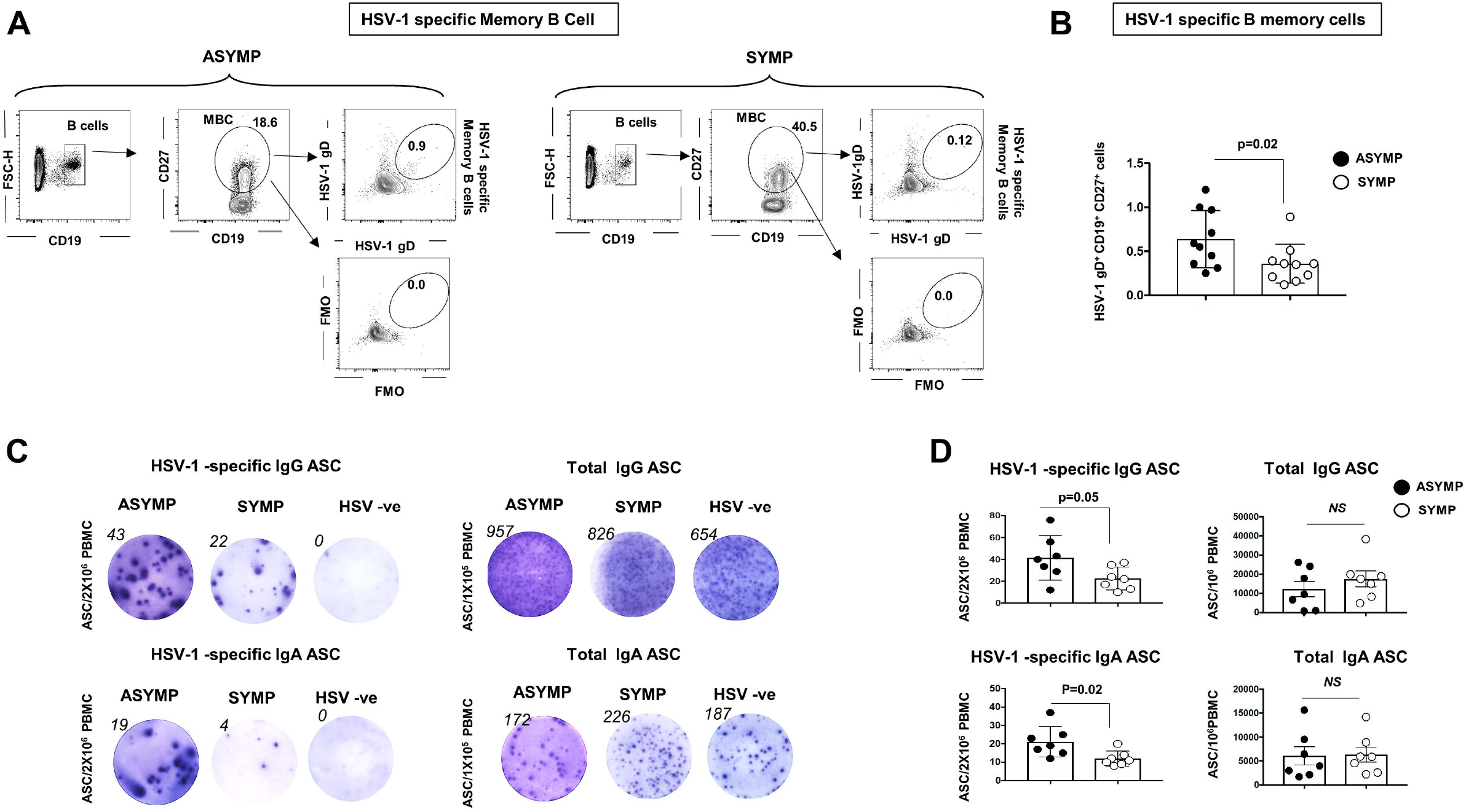
Circulating HSV-1 gD specific memory B cell profile in asymptomatic and symptomatic herpes infected individuals: PBMC from asymptomatic and symptomatic HSV-1 infected individuals were stained for HSV-1 gD antigen specific memory B cells. PBMC were also treated with (IL-2 & Resiquimod) for 5 days for polyclonal stimulation of memory B cells in to plasma cells (Antibody Secreting Cells-ASC). The stimulated cells were then analyzed for HSV-1 gD specific IgG/ IgA ASC and total IgG/ IgA ASC by ELISPOT. (**A**) FACS plot showing the gating strategy for HSV-1 specific memory B cells (CD19^+^CD27^+^ B cells) in PBMC of asymptomatic (ASYMP) (*Left panels*) and symptomatic (SYMP) (*Right panels*) HSV-1 infected individuals. (**B**) Graph showing percentage of HSV-1 specific memory B cells (CD19^+^CD27^+^ B cells) in PBMC of ASYMP and SYMP HSV-1 infected individuals. **(C)** Representative ELISPOT images for anti-HSV-1 IgG ASC (*Left, top panel*) anti-HSV-1 IgA ASC (*Left, bottom panel*) from PBMC of ASYMP, SYMP HSV-1 infected individuals and HSV-1 uninfected individuals (polyclonally stimulated for maturation of memory B cells to ASC). **(D)** Graph showing anti-HSV-1 IgG (*Left, top panel*) and IgA (*Left, bottom panel*) ASC; total IgG (*Right, top panel*) and total IgA (*Right, bottom panel*) ASC from PBMC of ASYMP, SYMP HSV-1 infected individuals and HSV-1 uninfected individuals. Statistical analysis was done using student’s *t* test. NS: not significant.

PBMCs from ASYMP, SYMP HSV-1 infected individuals and HSV-1 uninfected individuals were also treated with (IL-2 & Resiquimod) for 5 days for polyclonal stimulation of memory B cells into plasma cells (Antibody Secreting Cells-ASC). Stimulated cells were then analyzed for HSV-1 gD specific IgG/ IgA ASC and total IgG/ IgA ASC by ELISPOT. We detected an increased HSV-1 gD specific ASC (both IgG [41.36 ± 7.7 Vs 22.57 ± 3.92] and IgA [21.14 ± 3.15 Vs 12 ± 1.5] in ASYMP compared to SYMP (**Fig. 1C** and **1D)**. The ELISPOT findings confirm flow cytometry that ASYMP individuals have increased circulating memory cells binding to HSV-1 gD compared to SYMP individuals. As expected, HSV-1 and HSV-2 negative individuals showed no detectable HSV-1 gD specific IgG/ IgA ASC compared to ASYMP and SYMP herpes positive subjects (**Fig. 1C**).

### 2. Circulating HSV-1 specific switched memory B cells positively correlates with memory Tfh

CD19^+^CD27^+^IgD^−^ cells are known as “switched” memory B cells which indicates B-cell activation & development in germinal centers in lymph nodes or other secondary lymphoid tissues. PBMCs from asymptomatic and symptomatic HSV-1 infected individuals were stained for memory switched B cells (CD19^+^CD27^+^IgD^-^) and HSV-1 gD specific memory switched B cells (**Fig. 2A**). We found a trend towards increased switched memory B cells in ASYMP (0.771 ± 0.12%) compared to SYMP (0.345 ± 0.09%) (*P* = 0.04) (**Fig. 2B**), indicating that T cell dependent memory B cell is diminished during SYMP herpes infection.

**Figure 2:**
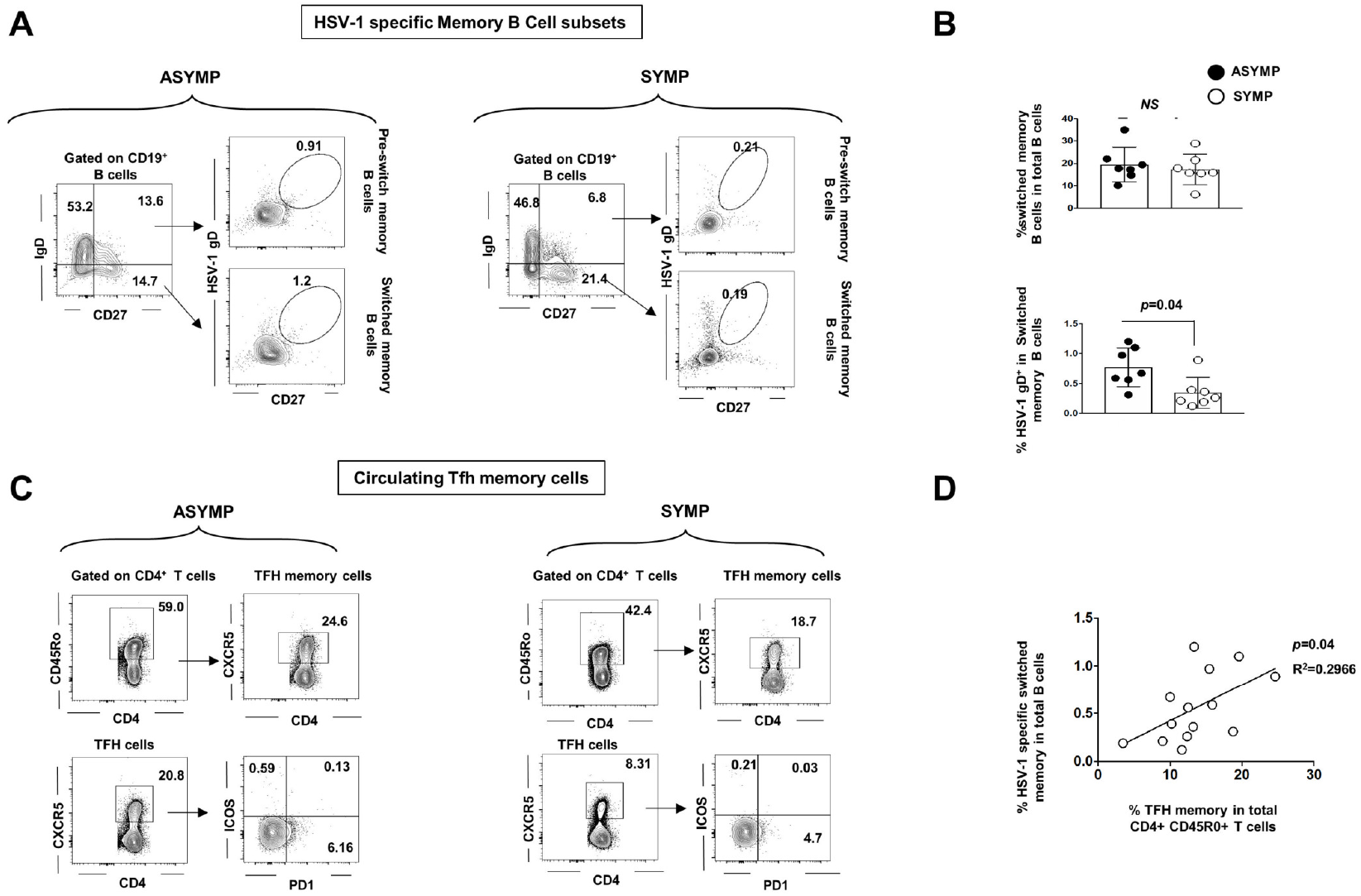
Correlation of HSV-1 specific memory B cell and memory T follicular helper T cells in herpes infected individuals: PBMC from asymptomatic and symptomatic HSV-1 infected individuals were stained for switched memory B cells (CD19^+^CD27^+^IgD^-^) and HSV-1 gD specific switched memory B cells. (**A**) FACS plot showing gating strategy for HSV-1 specific switched memory B cells (CD19^+^CD27^+^ B cells) in PBMC of asymptomatic (ASYMP) (*Left panels*) and symptomatic (SYMP) (*Right panels*) HSV-1 infected individuals. (**B**) Graph showing percentage of HSV-1 specific switched memory B cells (CD19^+^CD27^+^IgD^-^ B cells) in PBMC of ASYMP and SYMP HSV-1 infected individuals. (**C**) FACS plot showing gating strategy for T follicular helper cells (Tfh) (CD3^+^CD4^+^CXCR5^+^ T cells) and T follicular helper memory cells (Tfh memory) (CD3^+^CD4^+^CD45R0^+^CXCR5^+^ T cells) in PBMC of asymptomatic (ASYMP) (*Left panels*) and symptomatic (SYMP) (*Right panels*) HSV-1 infected individuals. (**B**) Graph showing correlation of the percentage of HSV-1 specific switched memory B cells (CD19^+^CD27^+^IgD^-^ B cells) and T follicular helper memory cells (Tfh memory) (CD3^+^CD4^+^CD45R0^+^CXCR5^+^ T cells) in PBMC of HSV-1 infected individuals. Statistical analysis was done using student’s *t* test. NS: not significant.

Switched memory B cells are derived from naïve B cells with T-cell help in extra-follicular or germinal centers. Hence, we explored the levels of circulating T follicular helper T memory cells (cTFH memory) that is found in circulation of asymptomatic and symptomatic HSV-1 infected individuals by flow cytometry. ASYMP (*n* = 7) and SYMP (*n* = 7) individuals PBMCs was also stained for Tfh cells (CD3^+^ CD4^+^ CXCR5^+^ T cells) and Tfh memory cells (CD3^+^CD4^+^CD45R0^+^CXCR5^+^ T cells) (**Fig. 2C**). We detected a positive correlation between the percentage of HSV-1 specific switched memory B cells (CD19^+^CD27^+^IgD^-^ B cells) and T follicular helper memory cells (Tfh memory) (CD3^+^CD4^+^CD45R0^+^CXCR5^+^ T cells) in PBMCs of HSV-1 infected individuals (*P* = 0.04).

### 3. HSV-1 binding antibody and neutralizing antibody levels in plasma of ASYMP and SYMP herpes infected individuals are similar

Circulating binding/neutralizing antibodies (secreted by ASC/plasma cells) are the first-line of B cell defense that target the virus. Serum from asymptomatic and symptomatic HSV-1-infected individuals were used in the estimation of anti-HSV-1 gD antibodies by ELISA. No significant difference in the anti-HSV-1 IgG and IgA antibody levels in plasma of ASYMP and SYMP herpes infected individuals was observed (**Fig. 3C**). Similarly, anti-HSV-1 neutralizing antibody titer of ASYMP (PRNT_50_ is 832) and SYMP (PRNT_50_ is 1024) herpes individuals did not vary significantly. PRNT50 represents reciprocal serum dilution at which 50% virus neutralization was observed.

**Figure 3:**
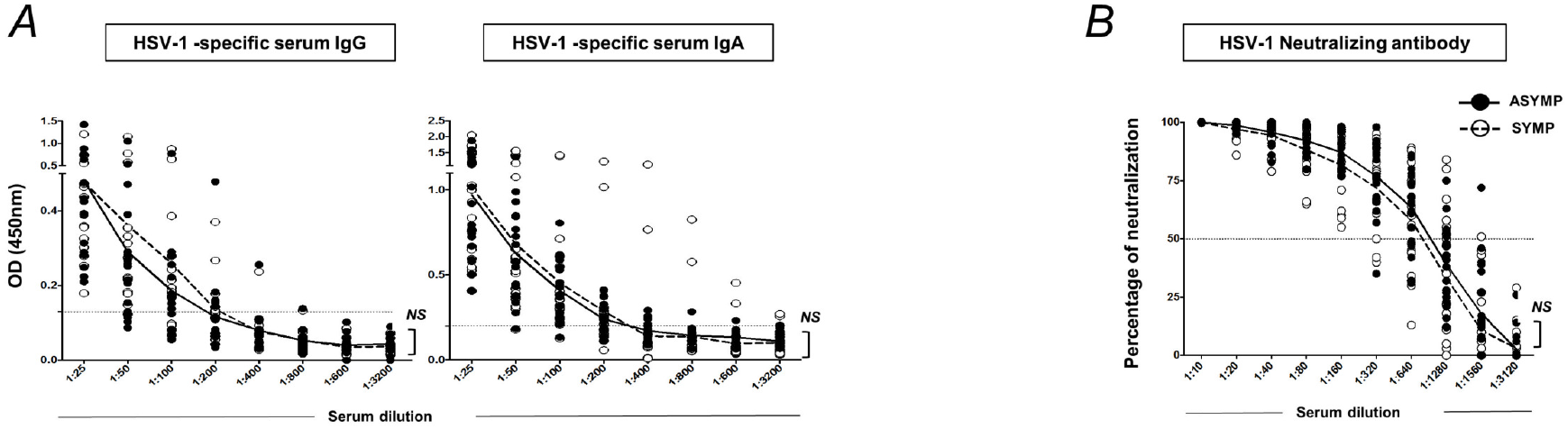
Anti-HSV-1 IgG and IgA antibody levels and neutralizing antibody titre in ASYMP and SYMP herpes infected individuals: Serum from asymptomatic and symptomatic HSV- 1 infected individuals were used for estimation of anti-HSV-1 gD antibodies by ELISA. (**A**) Graphs showing levels of anti-HSV-1 IgG antibody level (*left panel*) and anti-HSV-1 IgA antibody level (*left panel*) in serum of ASYMP and SYMP herpes infected individuals. (**A**) Graph showing anti-HSV-1 neutralizing antibody titer in serum of ASYMP and SYMP herpes infected individuals. Statistical analysis was done using student’s *t* test. NS: not significant.

### 4. Increased memory B cells in spleen of ASYMP HSV-1 reactivated mice

To further understand the role of circulating memory B cell in asymptomatic herpes, we used UV-B reactivation in the HSV-1 infected mouse model. Cornea of B6 mice were infected with HSV-1 McKrae (1×10^6^ pfu/eye) and virus reactivation was provoked at day 35 PI in latently infected mice, using a 60-second corneal UV-B irradiation. At day 6 post-reactivation, mice were categorized into ASYMP or SYMP depending on disease occurrence and euthanized and immune cells from peripheral blood, spleen and bone marrow were collected for flow cytometry staining of B memory cells. Memory B cells are B220^+^CD73^+^ B cells and its subsets B220^+^CD73^+^CD80^+^PD-L2^+^ B cells (**Fig. 4A**). The percentage of memory B cells B220^+^CD73^+^ in the spleen was observed to be higher in ASYMP (12.65 ± 0.55%) as compared to SYMP (7.64± 0.42%) while no difference was observed in the memory B cells bone marrow in ASYMP and SYMP infected mice (3.01± 0.03%Vs 2.35± 0.25%) (**Fig. 4B**).

**Figure 4:**
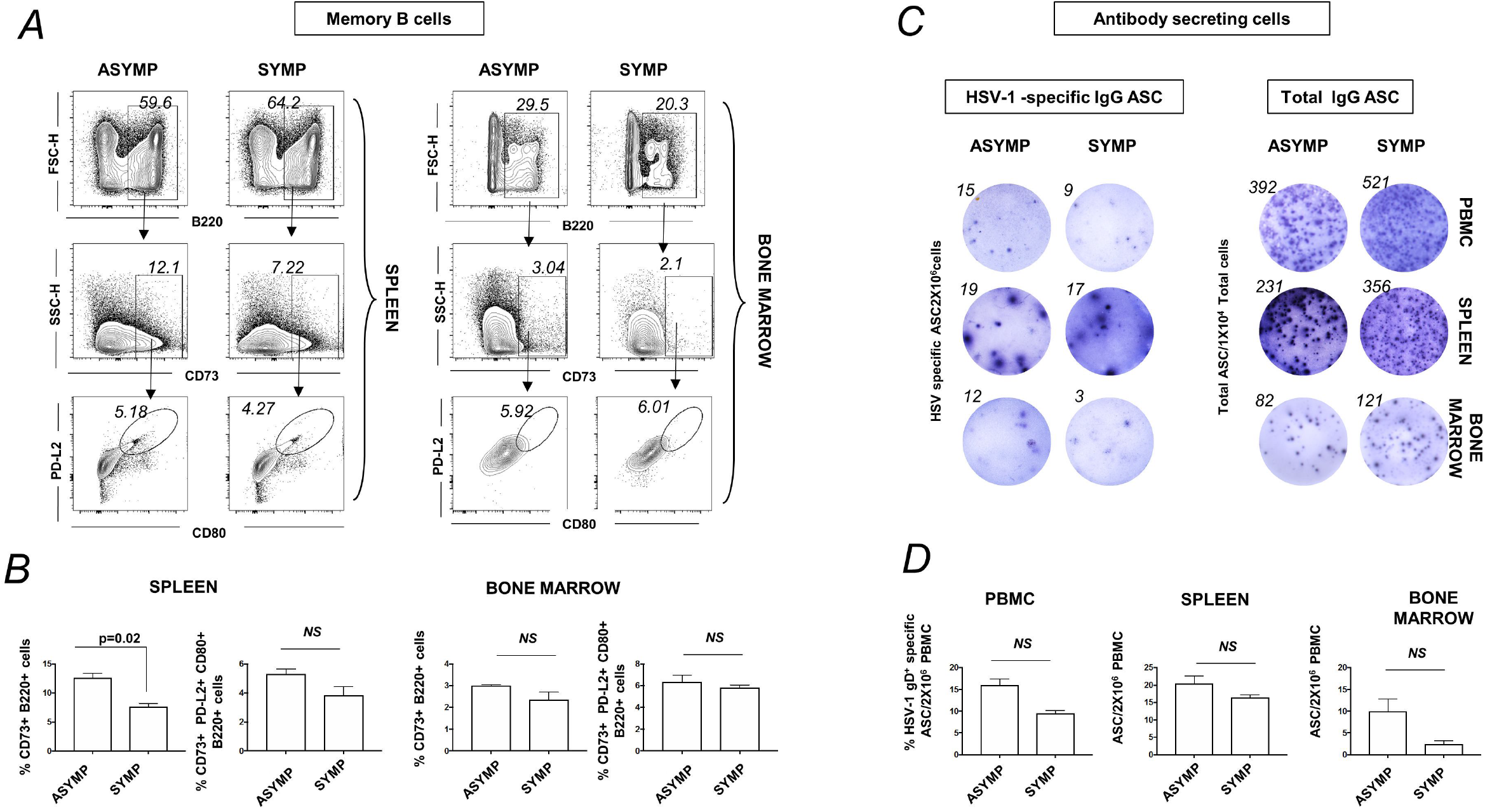
Memory B cell profile in PBMC, Spleen and bone marrow of ASYMP and SYMP HSV-1 reactivated mice: For this experiment, the cornea of B6 mice were infected with HSV-1 McKrae (1×10^6^ pfu/eye) by scarification and virus reactivation was provoked at day 35 PI in latently infected mice, using a 60 second corneal UV-B irradiation. At day 6 post-reactivation, mice were categorized into ASYMP or SYMP depending on disease occurrence. ASYMP and SYMP mice were euthanized and immune cells from peripheral blood, Spleen and bone marrow were collected for flowcytometry staining for memory B cells. (**A**) FACS plot showing representative plots for memory B cells (B220^+^ CD73^+^ B cells) and the subsets (B220^+^CD80^+^PD-L2^+^ B cells) in spleen (*Left panels*) and bone marrow (*Right panels*) of ASYMP and SYMP infected mice. (**B**) Graph showing percentage of for memory B cells (B220^+^CD73^+^ B cells) and the subsets (B220^+^CD73^+^CD80^+^PD-L2^+^ B cells) in Spleen (*left panel*) and bone marrow (*Right panel*) of ASYMP and SYMP infected mice. **(C)** Representative ELISPOT images for anti-HSV-1 IgG ASC (*Left panel*) and total IgG ASC (*Right panel*) from PBMC, spleen and bone marrow from ASYMP, SYMP infected mice (polyclonally stimulated with mouse polyclonal B cell activator (IL-2+Resiquimod) for maturation of memory B cells to ASC). Graph showing anti-HSV-1 IgG (*Left panel*) and total IgG (*Right panel*) ASC from PBMC of ASYMP, SYMP HSV-1 infected mice. Statistical analysis was done using student’s *t* test. NS: not significant.

Immune cells from peripheral blood, spleen and bone marrow were stimulated with mouse polyclonal B cell activator (IL-2 + Resiquimod) for maturation of memory B cells to ASC (using human B-Poly-S from Immunospot, OH, USA). The stimulated cells were incubated in ELISPOT plates to enumerate anti-HSV-1 IgG ASC and total IgG ASC (from PBMC, spleen and bone marrow). Anti-HSV-1 IgG ASC from PBMCs, spleen and BM depicted an increased trend in ASYMP compared to SYMP. The results from the mouse reactivation model confirm the results from humans showing an increased memory B cell profile in ASYMP herpes. However, we did not find any difference in the memory B cell frequency in the cornea of infected ASYMP mice compared to SYMP mice (data not shown).

### 5. Increased plasma cells in TG of ASYMP mice compared to SYMP mice

ASYMP and SYMP mice were euthanized and immune cells from peripheral blood, spleen, BM and TG cells were stained for plasma cells (CD138^+^ B cells) for flow cytometry. There was an increase in the percentage of plasma B cells (CD138^+^ B cells) in TG of ASYMP (5.23 ± 0.47%) as compared to SYMP (2.15 ± 0.50%) mice (**Fig. 5A**). ASYMP mice had increased HSV-1 specific plasma B cells (HSV-1 gD^+^ CD138^+^ B cells) (6.53 ± 0.15% Vs 4.54± 0.44%) compared to SYMP mice (**Fig. 5B**).

**Figure 5:**
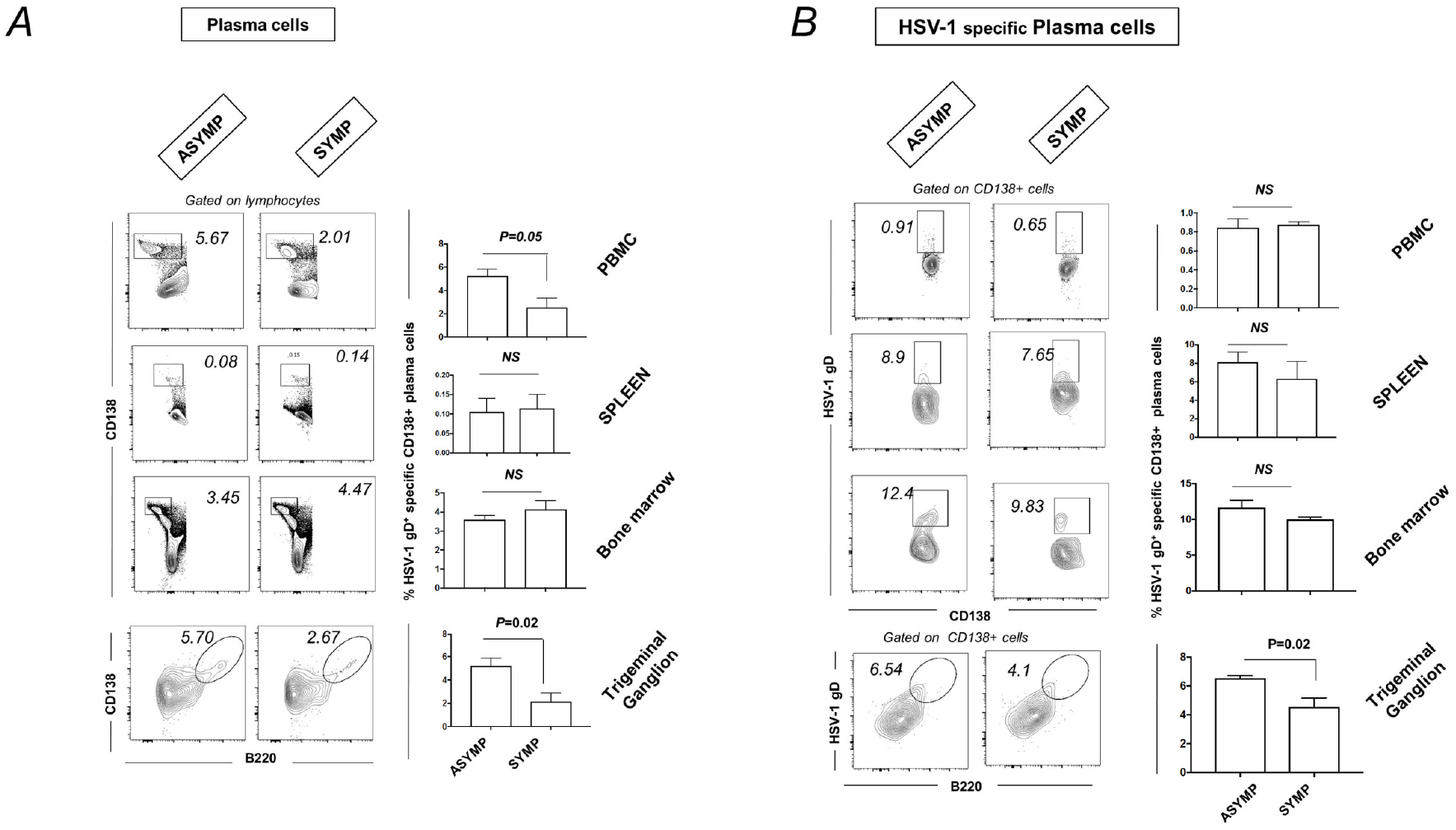
Plasma cell profile in PBMC, spleen, bone marrow and TG of ASYMP and SYMP HSV-1 infected mice: For this experiment, the cornea of B6 mice were infected with HSV-1 McKrae (1×10^6^ pfu/eye) by scarification and virus reactivation was provoked at day 35 PI in latently infected mice, using a 60 second corneal UV-B irradiation. At day 6 post-reactivation, mice were categorized into ASYMP or SYMP depending on disease occurrence. ASYMP and SYMP mice were euthanized and immune cells from peripheral blood, Spleen and bone marrow were collected for flowcytometry staining for plasma cells. (**A**) Representative plots for plasma B cells (CD138^+^ B cells) in PBMC, Spleen, PBMC and TG (*Left panels*) of ASYMP and SYMP mice is shown. Graph showing percentage of plasma B cells (CD138^+^ B cells) in PBMC, Spleen, PBMC and TG (*Right panels*) of ASYMP and SYMP mice. **(B**) Representative FACS plot showing HSV-1 specific plasma B cells (HSV-1gD^+^CD138^+^ B cells) in PBMC, Spleen, PBMC, and TG (*Left panels*) of ASYMP and SYMP mice. Graph showing percentage of HSV-1 specific plasma B cells (HSV-1gD^+^CD138^+^ B cells) in PBMC, Spleen, PBMC and TG (*Right panels*) of ASYMP and SYMP mice.

### 6. Tfh and Tfh memory cell profile in spleen of ASYMP and SYMP HSV-1 infected mice

Spleen cells were collected for flow cytometry staining of T follicular helper (Tfh) (CD3^+^CD4^+^CXCR5^+^PD1^+^ cells) and T follicular helper memory (Tfh memory cells) (CD3^+^CD4^+^CD44^+^CXCR5^+^ PD-1^+^ cells). There was no difference detected in the percentage of Tfh cells (top) Tfh memory (bottom) in spleen of ASYMP (Tfh: 11.7± 1.2%, Tfh memory: 7.45 ± 0.45%) and SYMP (Tfh: 12.57± 0.88%, Tfh memory: 8.99 ± 0.32%) infected mice (**Figs. 6A** and **6B**).

**Figure 6.**
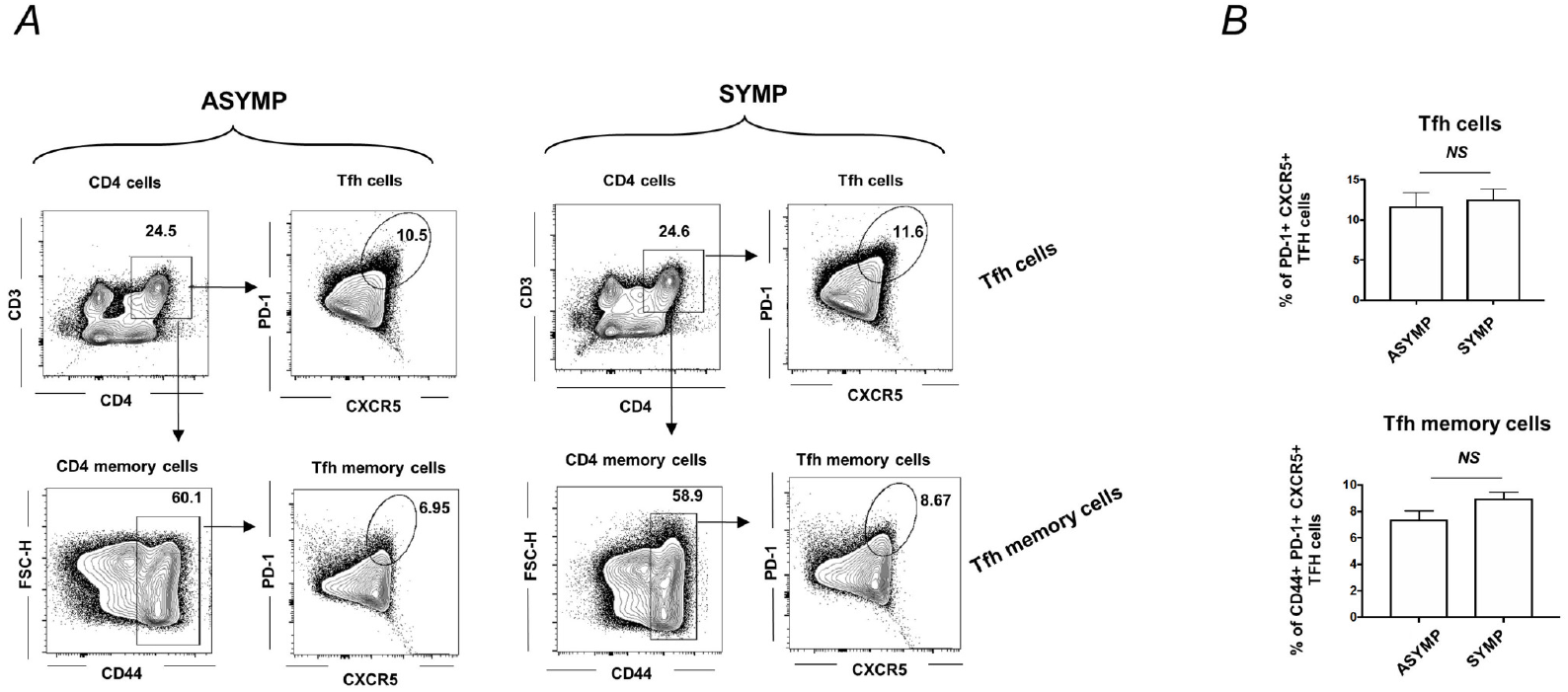
T follicular helper (Tfh) and T follicular helper memory (Tfh memory) cell profile in spleen of ASYMP and SYMP HSV-1 infected mice: B6 mice were infected with HSV-1 McKrae (1×10^6^ pfu/eye) by scarification and at day 35 PI, reactivation was done by a 60 second corneal UV-B irradiation. Mice were categorized as ASYMP /SYMP and euthanized at day6 post-reactivation. Spleen cells were collected for flowcytometry staining of T follicular helper (Tfh) (CD3^+^CD4^+^CXCR5^+^PD-1^+^ cells) and T follicular helper memory (Tfh memory) (CD3^+^CD4^+^CD44^+^CXCR5^+^PD-1^+^ cells). (**A**) Representative FACS plots for Tfh cells (CD3^+^CD4^+^CXCR5^+^PD-1^+^ cells) is shown in *top panel* and Tfh memory cells (CD3^+^CD4^+^CD44^+^CXCR5^+^PD-1^+^ cells) is shown in bottom panels for ASYMP (*left panel*) and SYMP (right panel) infected mice. (**A**) Graph showing percentage of Tfh cells (top) Tfh memory (bottom) in spleen of ASYMP and SYMP infected mice is shown.

**Figure 7.**
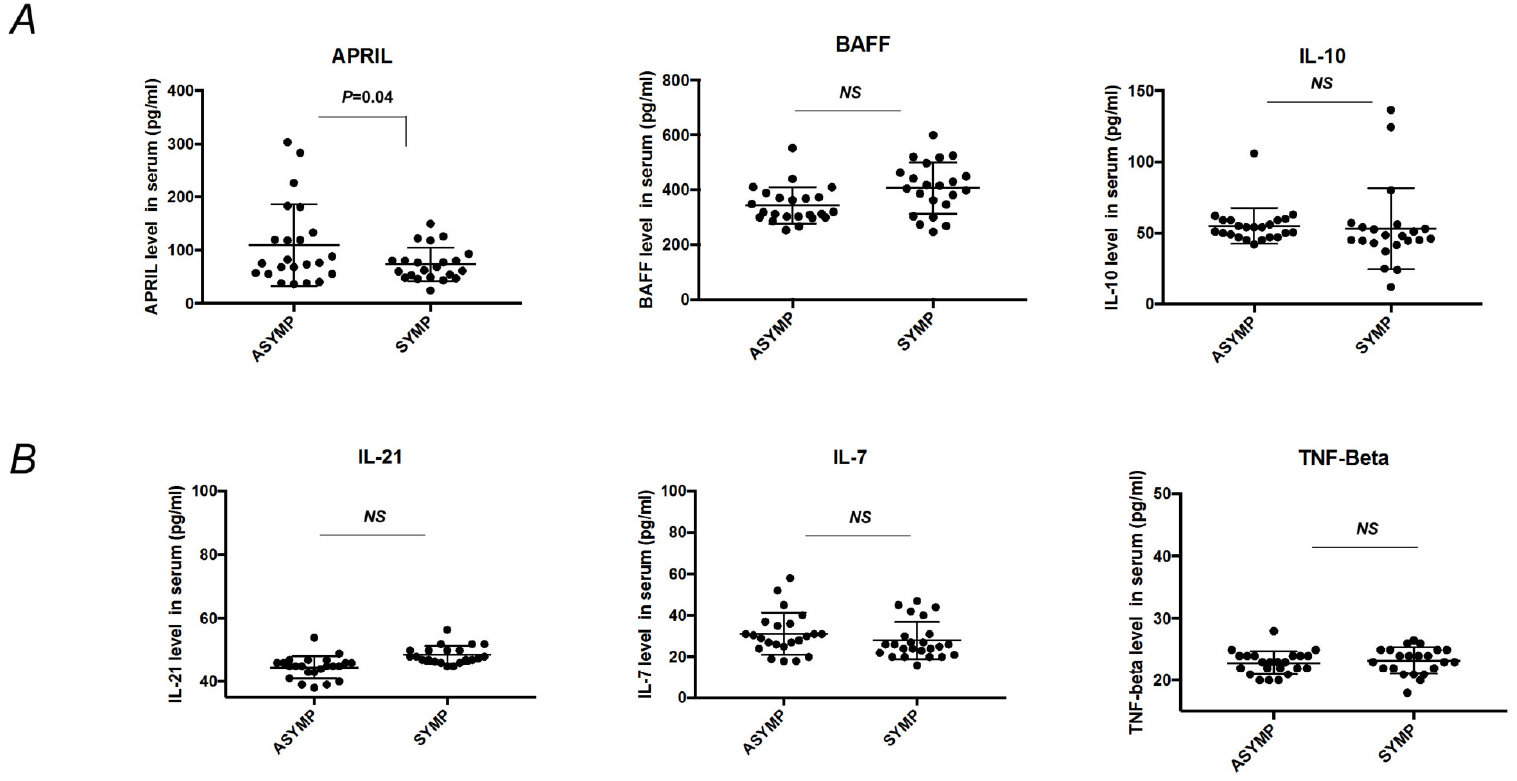
B cell ligand and cytokine level in serum of asymptomatic ans symptomatic herpes: Asymptomatic (n=20) and symptomatic (n=20) patients’ serum were assayed for cytokines involved in B cell development. Graph showing serum level of A). APRIL (B cell proliferation inducing ligand), BAFF (B cell activating factor), IL-10 (B regulatory cell cytokine) B). IL-21 (involved in expansion and differentiation of plasma cells), IL-7 (B cell development cytokine) and TNF-β (B cell development cytokine) by Luminex. Statistical analysis was done using student’s *t* test. NS: not significant.

### 7. Increased APRIL (B cell proliferation inducing ligand) in asymptomatic vs symptomatic herpes individuals

Asymptomatic and symptomatic patients’ serum (heat-inactivated at 56°C for 30 minutes) were assayed for cytokines involved in B cell development APRIL (a B cell proliferation inducing ligand), BAFF (B cell activating factor), IL-10, IL-21, IL-7 and TNF-β (B cell development cytokine) by Luminex. We found increased APRIL levels in serum of ASMP herpes individuals (109.4 ± 16.03 pg/ml) compared to SYMP herpes individuals (73.5 ± 6.7 pg/ml). The levels of other B cell associated cytokines such as BAFF, IL-10, IL-21, IL-7 and TNF-β did not vary significantly between asymptomatic and symptomatic herpes individuals.

## DISCUSSION

The development of an effective therapy to alleviate symptoms of recurrent Herpes Stromal Keratitis is dependent on our understanding of immune response to herpes infection. In the current study, we are examining the role of B cell mediated immunity in the human response to HSV-1 reactivation. Anti-herpes antibodies are reported to not be protective during primary herpes infection (32). Reports are indicating that high titers of herpes neutralizing antibodies in the blood of symptomatic herpes is neither associated with the frequency of symptomatic herpes (30, 31) nor protection after vaccination (32, 33). Thus, the general understanding is that anti-herpes antibodies although are neutralizing or binding, are not protective (34). Another theory states that certain HSV-specific antibodies may function protectively in tissue near the site of viral release. However, circulating antibody levels do not reflect this tissue-based event (35). Cell-to-cell spread of the virus is sufficient for propagation and development of symptomatic reactivation even in the presence of effective neutralizing antibodies. Therefore, the concentration of such antibodies is below the threshold of efficacy (36). Recent evidence suggests a more migratory role of B cells to the site of herpes reactivation in skin (15).

Memory B cells can survive for long periods & can induce faster and stronger humoral responses when they reencounter the same antigen (37), in contrast to plasma cells which provide the first line of protection against infection but do not respond to the second infection because of low expression of membrane-bound Ig (38). There is evidence which indicates that human memory B cells reside mostly in the spleen, & some memory B cells recirculate in the blood. In the present study, we evaluated whether and if memory B cell response is associated with protection in HSV-1 infection, a question that is not yet fully understood. The frequency of memory B cells and a counterpart of follicular helper T cells (Tfh) found in circulation of asymptomatic and symptomatic HSV-1 infected individuals were studied by flow cytometry. The memory B cells can respond to re-infection with rapid formation of extra-follicular foci, thereby broadening the oligo-clonal repertoire of germ-line encoded B cells causing a broadening of the antiviral B-cell repertoires (37). Our results show that asymptomatic HSV-1 infected individuals do not only have an increased frequency of circulating memory B cell specific to HSV-1 gD antigen but also an increased HSV-1 specific memory B cell functional response as estimated by ELISPOT and the levels of circulating HSV-1 specific memory B cells were directly proportional to the level of circulating follicular helper CD4 T cells. We also examined if there were any differences in B cell development associated cytokines and ligands (APRIL, BAFF, IL-10, IL-21, TNF-β)□ in the serum of ASYMP and SYMP herpes individuals. We found that APRIL, released by myeloid and stromal cells, which can enhance the longevity of humoral immune response, was increased in the serum of asymptomatic herpes individuals.

Currently, there is no effective cure or vaccine against HSV infection. HSV vaccines from the past focused on subunit formulations designed to elicit neutralizing antibodies targeting the envelope glycoprotein D (gD). Passive antibody transfer and sequential infection experiments demonstrated ‘original antigenic suppression’, a phenomenon in which antibodies suppress memory responses to the priming antigenic site. While these vaccines elicited high levels of neutralizing antibodies in animals and humans, they failed to protect against HSV-2 infections in clinical trials. Our observation showed that in spite of an increased memory B cells in circulation, the herpes antibody levels remained the same between ASYMP and SYMP. Thus, we wanted to explore if HSV-1 specific memory B cells found in circulation would possibly have any role or effect that is confined locally at the site of reactivation in asymptomatic herpes using UV-B-induced HSV-1 reactivation mouse model. Although the spleen and tonsil are the major reservoirs for antigen-specific human memory B cells, they appear to be dispensable for preserving immunological memory following a re-encounter with the antigen (38). There was a trend towards an increased memory B cell response, but not the antibody-secreting plasma cells in PBMCs, spleen and bone-marrow of asymptomatic compared to symptomatic HSV-1 reactivated mice. In another recent study, investigators have shown that maternal antibodies were found at fetal trigeminal ganglia conferring complete protection for newborn mice against HSV infection. In our study, we detected an increase in HSV-1 specific plasma B cells in the TG of asymptomatic mice as compared to symptomatic mice. Thus, the discordance between HSV-specific memory B cells and antibody levels in circulation in human natural infection can be explained by a virus driven memory B cell recruitment mechanism that leads to antibody production at the site of reactivation (TG) using mouse model. Our findings suggest that circulating HSV-specific antibody-producing memory B cells recruited locally at the TG site could contribute to protection from symptomatic recurrent ocular herpes.

## ACKNOWLEDGEMENTS

This work is supported by a grant from Trefoil Therapeutics, Inc. and by Public Health Service research grants EY019896, EY14900 and EY024618 from the National Eye Institutes (NEI) and AI150091, AI143348, AI147499, AI143326, AI138764, AI124911 and AI110902 from the National Institutes of Allergy and Infectious Diseases (NIAID) to LBM and from The Discovery Center for Eye Research, and in part by Research to Prevent Blindness.

